# Prediction of late blight severity in a large panel of potato genotypes using low-altitude aerial images and machine learning methods

**DOI:** 10.64898/2026.04.06.716456

**Authors:** Hildo Loayza, Johan Ninanya, Susan Palacios, Luis Silva, Fernando Pujaico Rivera, Javier Rinza, Manuel Gastelo, Mariela Aponte, Jan F. Kreuze, Hannele Lindqvist-Kreuze, Bettina Heider, Moctar Kante, David A. Ramírez

## Abstract

Potato (*Solanum tuberosum* L.) is a staple crop crucial to global food security, yet its production is severely threatened by late blight (LB), caused by *Phytophthora infestans*, one of the most destructive plant diseases worldwide. Breeding programs for LB resistance have traditionally relied on labor-intensive and subjective visual assessments, which limit scalability and consistency, particularly in early-generation trials. Unmanned aerial vehicle (UAV)-based remote sensing combined with machine learning (ML) offers a promising alternative for objective, high-throughput disease phenotyping. This study evaluated the potential of UAV-derived multispectral imagery and ML techniques to estimate LB severity across large and genetically diverse potato breeding populations, comprising 2,745 clones in one trial and 492 accessions in another, conducted in Oxapampa, Pasco, Peru. We compared vegetation index–based approaches with a machine learning framework that integrates K-means clustering and Kernel Ridge Regression (KRR) and assessed their ability to capture genotypic variation and support selection decisions. NDVI consistently showed a strong correlation with visually assessed LB severity, particularly at advanced stages of disease development, enabling objective discrimination between healthy and diseased canopy tissues. However, the KRR-based approach outperformed linear NDVI-based models by capturing nonlinear relationships between spectral responses and disease progression. Estimates of LB severity derived from NDVI and KRR models, expressed as best linear unbiased estimates (BLUEs), showed strong and biologically consistent relationships with the area under the disease progress curve (AUDPC), particularly during later UAV acquisitions. Selection coincidence between UAV-derived estimates and AUDPC-based rankings was substantially higher at intermediate to advanced stages of disease progression, suggesting that UAV assessments at these stages may capture sufficient phenotypic variation to distinguish genotypes. These findings indicate that UAV-based multispectral phenotyping, especially when integrated with ML, provides a practical and scalable approach for assessing LB severity in potato breeding programs while reducing the need for time-consuming field evaluations.

## Introduction

Potato (*Solanum tuberosum* L.) is a staple crop of critical importance to global food security and is the most widely grown non-cereal crop worldwide [1,2]. Potatoes contribute significantly to caloric intake, nutrition, and rural income [3]. However, the constant threat of numerous diseases, particularly late blight (LB), poses a severe challenge to potato production [4,5]. The pathogen *Phytophthora infestans* (Mont.) de Bary is an oomycete that causes LB, a highly virulent disease that can devastate potato crops. Early-season infections can lead to significant yield losses of up to 80%. This results in substantial economic hardship for farmers who rely on this crucial crop, with annual losses estimated to be between 3 and 10 billion dollars [4,6,7]. The environmental, health, and financial implications of LB management further underscore the need for effective, sustainable solutions. LB spreads rapidly under optimum environmental conditions, especially at temperatures ranging between 10 and 23 °C and a relative humidity of more than 80% for 10 h daily [8–10]. It causes damage to foliage and tubers, which can lead to the loss of the entire crop within one week [11].

Potato breeding programs play an important role in LB integrated management by developing varieties with disease resistance, hence reducing reliance on chemical controls, lowering production costs, and contributing to sustainable agriculture [12–14]. The breeding process typically relies on visual disease assessments under controlled infection conditions, in which trained experts evaluate disease progression across several genotypes [15,16]. These assessments involved estimating the percentage of infected leaf area per plant, assuming that LB was the only foliar disease present in the experiment. However, this method is inherently labor-intensive, time-consuming, and subject to observer bias, which limits its scalability and precision [17]. Furthermore, breeding programs targeting LB resistance face numerous challenges, including the need to test thousands of genotypes in early generation trials [18]. This large-scale phenotyping requires considerable human resources and incurs high costs due to the intensive, repeated manual assessments conducted throughout the growing season. Additionally, environmental variability and subjective assessment practices can introduce inconsistencies, complicating the accurate identification of resistant genotypes in different environments. These challenges underscore the need for alternative phenotyping approaches to increase consistency, scalability, and cost efficiency [19].

Advances in remote sensing offer potential solutions to these limitations in phenotyping. Remote sensing tools, including sensors mounted on unmanned aerial vehicles (UAVs), with improved spatial and temporal resolutions, now enable high-throughput, non-destructive data collection across extensive breeding plots [20,21]. These tools provide rapid and objective data on disease progression, enabling breeders to efficiently and at scale assess crop health. For example, RGB and multispectral images can be used to compute vegetation indices that are highly correlated with plant health indicators, providing an alternative approach to disease assessment [22]. The combination of multispectral imagery with machine learning (ML) and deep learning (DL) models enables the detection of complex disease-related patterns in large datasets, facilitating efficient disease detection and classification [17,23,24]. Thus, multispectral images acquired from UAVs, when integrated with ML algorithms, have become powerful tools for the classification, segmentation, and detection of plant diseases [17,23]. In potato disease phenotyping, particularly for LB, UAV-based multispectral imaging and ML models have demonstrated promising results [17,25]. Despite these advancements, applying such tools to early-generation trials with several thousand genotypes remains challenging due to data-processing requirements and the need for consistent, reliable phenotyping.

This study aimed to evaluate the potential of UAV-based imagery, along with vegetation index-based approaches and ML techniques, to assess LB severity across a large panel of early-generation potato breeding trials. Specifically, the study’s objectives are as follows: i) to quantify the extent to which aerial phenotyping captures genetic variation in LB resistance, by comparing image-derived indicators with conventional field assessments across diverse genotypes; ii) to analyze the potential reduction of temporal assessments when UAV-based imagery is used to monitor genotypes’ LB severity. We hypothesized that ML modeling can effectively capture nonlinear relationships in multispectral reflectance data obtained from aerial imaging and disease severity. This capability may enhance the screening process for LB resistance across a large panel of genotypes. As a result, ML could reduce the time required for numerous field assessments typically conducted through conventional phenotyping, especially during critical developmental stages.

## Materials and methods

### Study area

Two potato trials were conducted in Oxapampa, Pasco, Peru (10.6039° S, 75.4153° W, 1,827 m a.s.l.), with the first trial (Trial A) from September 24, 2019, to January 7, 2020, and the second trial (Trial B) from October 21, 2019, to February 11, 2020 (Fig 1). Oxapampa is a site with high LB disease pressure [26], characterized by a temperate, rainy, and very humid climate year-round, with minimum and maximum temperatures ranging from 11 to 17 °C and 25 to 29 °C, respectively, and annual precipitation between 1200 and 3000 mm [27]. In both trials, the minimum and maximum temperatures ranged from 8.6 to 16.2 °C and 18.9 to 27.0 °C, respectively, whereas the relative humidity ranged from 69.0 to 94.7% (S1 Fig). During the crop growing season, the total precipitation was 559.6 mm for Trial A and 894.7 mm for Trial B (S1 Fig). The soil had a loam texture (46, 37, and 17% of sand, silt, and clay, respectively) with phosphorus, potassium, and organic matter contents of 53.1 ppm, 235 ppm, and 4.55%, respectively, as determined by the Soil, Plant, Water, and Fertilizer Analysis Laboratory of the National Agrarian University-La Molina, Lima, Peru.

**Fig 1.**
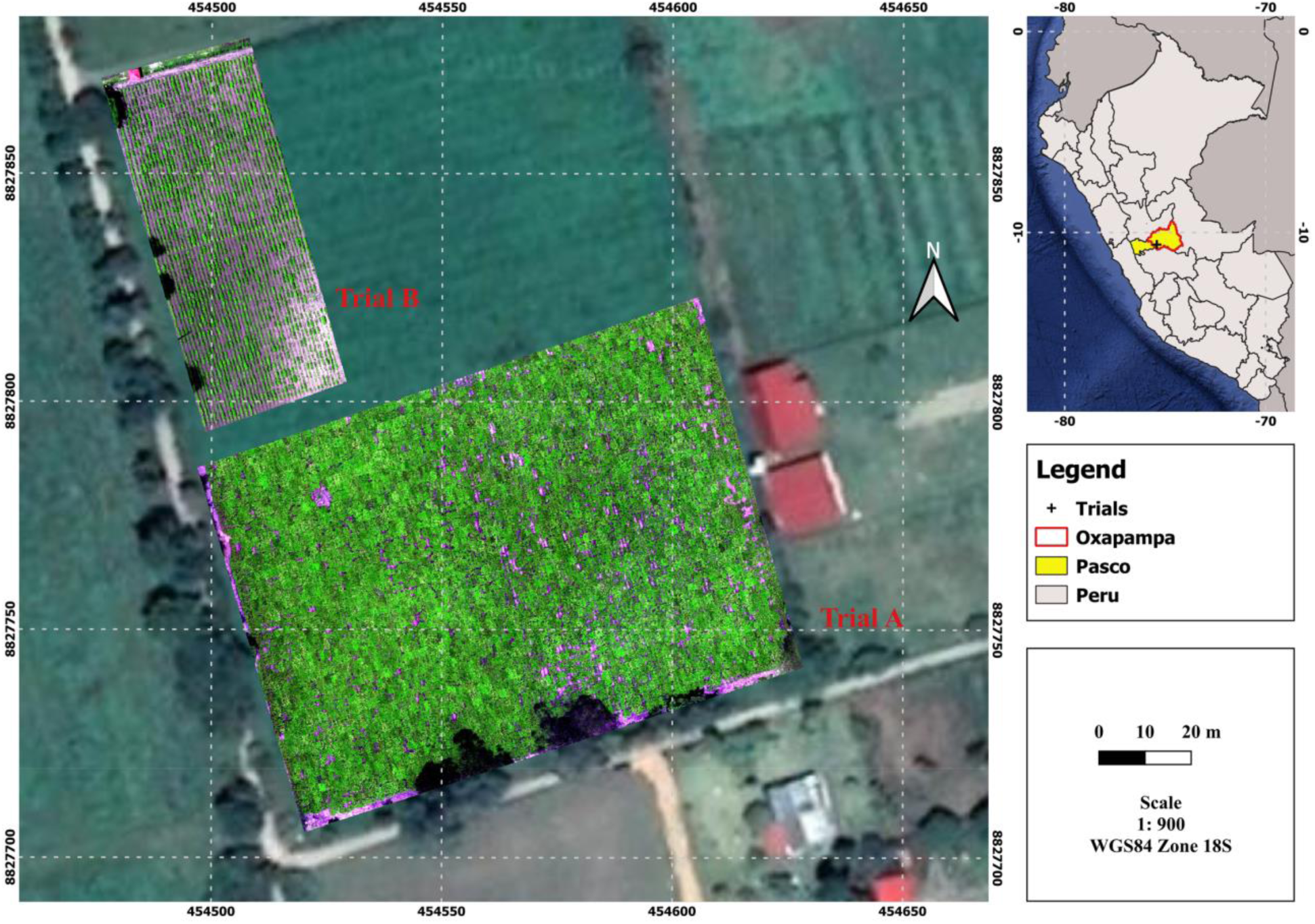
The study area. Location of the study area showing two potato trials in Oxapampa, Pasco, Peru: one including 2,745 clones with two replications over 0.9 ha (Trial A), and another involving 492 accessions with three replications covering 0.3 ha (Trial B).

**Fig 2.**
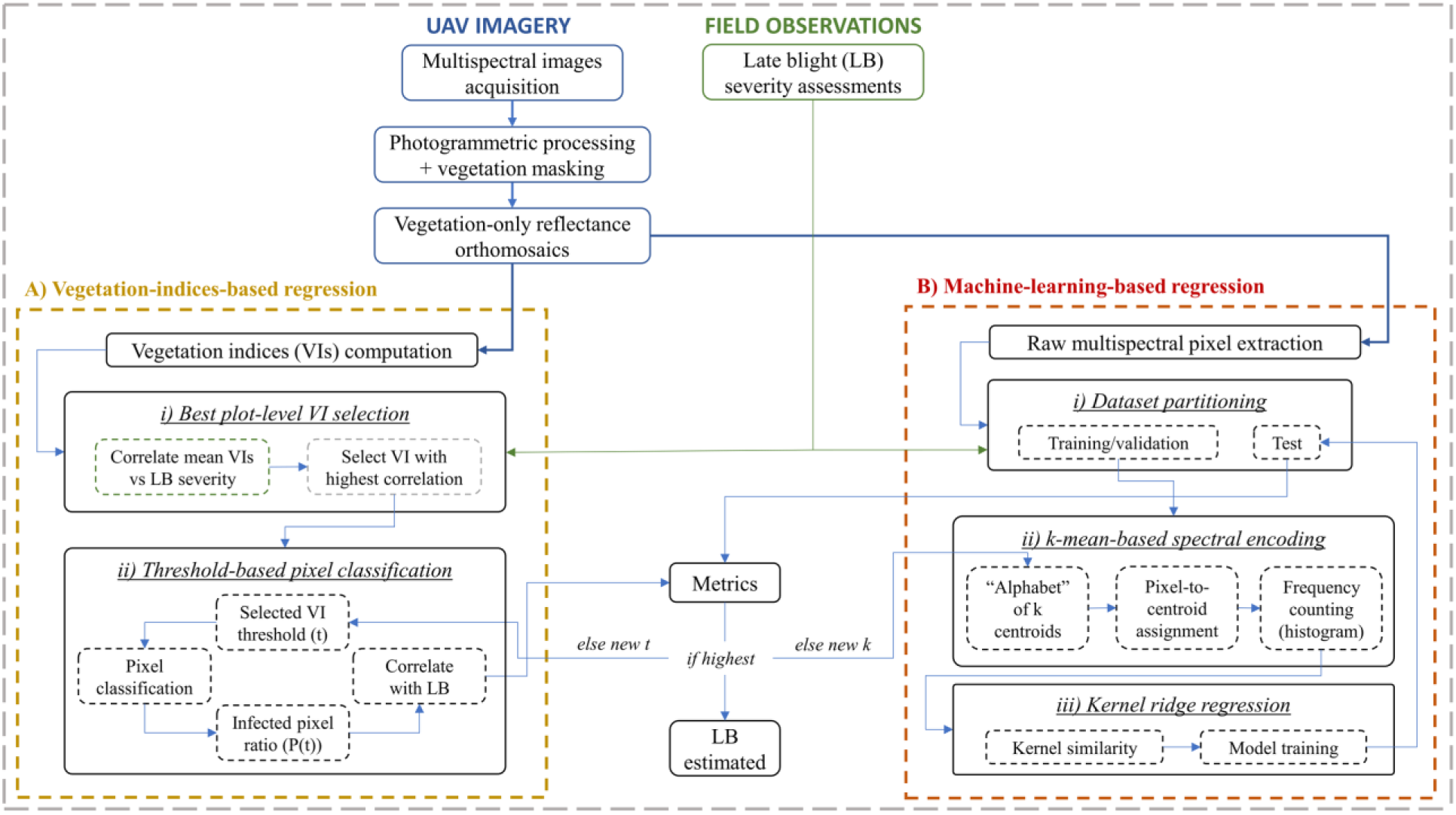
Workflow of NDVI- and KRR-based LB severity estimation. Methodology workflow of the two approaches used in this study: (A) Estimation of late blight (LB) severity using linear regression based on vegetation indices, and (B) a regression-based LB severity prediction system using Kernel Ridge Regression (KRR) combined with K-means clustering. In approach B, a dataset composed of five-dimensional multispectral pixel samples is first processed using K-means clustering with *K* centroids. This procedure generates a histogram vector with *K* elements, which is then used as input to the KRR model to estimate LB severity.

### Plant material and experimental design

In Trial A, 2,745 potato clones were evaluated as part of the preliminary yield trials of cycle 2 of the CIP LB-resistant and heat-tolerant (LBHT) breeding pipeline using a 43×65 resolvable row-column experimental design with two replications. In Trial B, a heterogeneous set of 492 potato accessions, comprising both diploid and tetraploid accessions from the CIP genebank core collection, was planted in a 12×41 row-column experimental design with three replications. In both trials, the plots consisted of single rows with five plants each, with a spacing of 0.3 m between plants and 0.9 m between plots. A common set of three varieties with known responses to LB—Amarilis (moderately resistant), Yungay (highly susceptible), and Kory (resistant)—was included in both trials as controls [15]. In Trial A, Amarilis and Yungay were replicated 34 times, whereas Kory was replicated 32 times. In Trial B, all three varieties were replicated 13 times each. Moreover, Yungay and Amarilis were planted around the field as “spreader rows” to allow for inoculum establishment and dissemination [28]. A description of the potato clones from Trials A and B is available in the CIP Dataverse repository [29,30].

Both trials were conducted according to standard agronomic practices [15] until all plants had emerged and grown sufficiently large for disease assessment, which was performed 40 days after planting (DAP). Consequently, to prevent early LB infection and before allowing natural LB infestation, fungicide Dithane M-45 WP NT (Mancozeb, BASF®, Peru) was applied preventively twice, at 80% and at full emergence. In both trials, fertilizers were applied at sowing at a rate of 200-180-160 kg ha^−1^ of N-P_2_O_5_-K_2_O, supplied as ammonium nitrate (238.3 kg), diammonium phosphate (373.5 kg), and potassium sulfate (305.4 kg). Additionally, 238.3 kg of ammonium nitrate was applied at 30 DAP (first hilling) to complete the remaining 50% of the required nitrogen. Potatoes were grown under rainfed conditions (S1 Fig), with total precipitation exceeding the 500 mm crop water requirement for 120-day potato crops [31].

### Field late blight severity assessment, image acquisition, and preprocessing

LB severity was assessed as the percentage of LB-infected foliar area per plant, visually estimated by trained experts assuming LB was the only foliar disease present in the plant, and recorded on a 0–100% scale in 5% increments [15]. LB severity assessments were conducted weekly at the plot level, comprising seven assessments in Trial A (starting at 41 DAP) and five assessments in Trial B (starting at 38 DAP). From these sequential assessments, disease development over time was quantified by calculating the Area Under the Disease Progress Curve (AUDPC) using the trapezoidal integration method [15,32].

Two UAV flights were conducted to acquire multispectral aerial images corresponding to two LB severity assessments: 64 and 86 DAP for Trial A, and 38 and 58 DAP for Trial B, respectively. These flights generated four datasets by combining LB severity observations with corresponding multispectral imagery. A RedEdge M camera (MicaSense, Seattle, USA) was mounted on a DJI Inspire 2 quadcopter (Shenzhen, China) to capture 16-bit images in five spectral bands: blue (475 ± 20 nm), green (560 ± 20 nm), red (668 ± 10 nm), red edge (717 ± 10 nm), and near-infrared (NIR; 840 ± 40 nm). The UAV flight plan was designed to ensure 75% overlap between images in both the frontal and lateral directions. Images were acquired at altitudes of 50 m in Trial A and 40 m in Trial B, resulting in spatial resolutions of 3.4 × 3.4 cm^2^ and 2.9 × 2.9 cm^2^, respectively. Flights occurred between 11:00 and 14:00 h local time, when potato plants exhibit maximum non-photochemical quenching [33]. Images of a calibrated reference panel were acquired before and after each flight to convert the radiance measurements into reflectance values. The multispectral dataset was processed in Pix4Dmapper (Prilly, Switzerland) for radiometric correction, geometric alignment, and mosaicking, producing a five-band reflectance orthomosaic for each flight in both trials.

### Late blight severity estimation

As a first step, reflectance orthomosaics from both trials were segmented using the Dzetsaka plug-in for QGIS, which implements a K-Nearest Neighbors (KNN) classifier [34]. The classifier was trained to label pixels and generate soil masks for each assessment date and trial; these masks were subsequently applied to produce orthomosaics containing only vegetation pixels from the potato plants. Subsequently, polygonal grids were created, labeled, and assigned to each plot according to the experimental design, resulting in 5,790 grids in Trial A and 1,476 grids in Trial B per dataset. Two methodologically distinct approaches were implemented to estimate LB severity (Fig 2): i) a vegetation-index-based linear regression approach applied at both the plot and pixel levels; and ii) a machine-learning-based approach that combines K-means clustering and Kernel Ridge Regression (KRR) using multispectral reflectance data.

**Fig 3.**
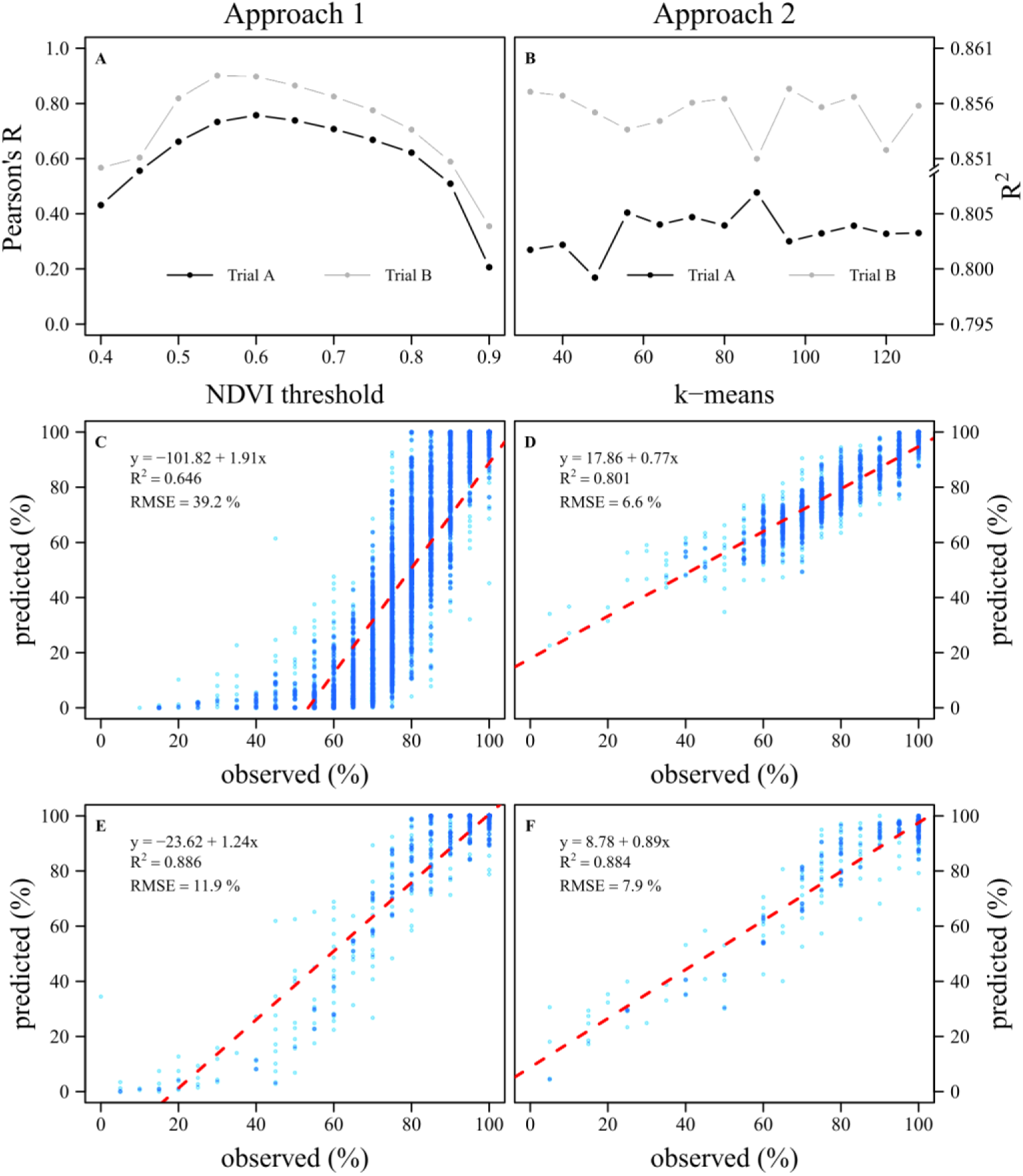
Optimization and validation of NDVI and KRR-based models for LB severity. (A) Pearson correlation coefficient (R) between observed and predicted Late Blight (LB) severity across a range of NDVI thresholds for Trials A and B. (B) Selection of the optimal number of clusters (*K*) in the K-means algorithm based on R^2^ values from linear regressions between observed and predicted LB severity using Gaussian kernel ridge regression (KRR) models during the training and validation stages for Trials A and B. (C) Linear regression between observed and predicted LB severity using the optimal NDVI threshold for Trial A (2,745 potato clones). (D) Linear regression between observed and predicted LB severity using a Gaussian KRR model applied to the testing subset and the optimal K value for Trial A. (E) Same as panel C, but for Trial B (492 potato accessions). (F) Same as panel D but for Trial B. All analyses were conducted using data acquired at 86 Days After Planting (DAP) for Trial A and 58 DAP for Trial B.

#### First approach: vegetation-index-based regression

This approach comprised two key steps (Fig 2A): i) identifying the vegetation index that best represents LB severity assessments, and ii) classifying potato plant cover as LB-infected or healthy using a threshold criterion. First, the plot-level mean values of four widely used vegetation indices— the Normalized Difference Vegetation Index (NDVI; [35]), Normalized Difference Red Edge (NDRE; [38]), Red Edge reflected band (RE), and Green Normalized Difference Vegetation Index (GNDVI; [39])—were computed for each LB severity observation dataset. These mean values were correlated with their corresponding LB severity assessments, and the vegetation index with the highest Pearson correlation coefficient (R) was selected as the “best” index for representing LB severity. Second, polygonal grids were used to crop the selected vegetation index images for each dataset. The resulting grid-based images were processed using a custom object-detection algorithm developed in MATLAB [40]. This algorithm identifies clusters of pixels in each grid (plot), encloses them within rectangular regions, labels them, and extracts all the associated pixel values [41]. Subsequently, for each plot, the full range of vegetation index values (0–1) was evaluated in increments of 0.05. Each increment was used as a threshold to classify the pixels as either LB-infected (pixel value ≤ threshold) or healthy (pixel value > threshold). For each threshold, the ratio of LB-infected pixels to the total number of pixels within each polygonal grid was determined. These ratios were compared with their corresponding LB severity assessments, and the threshold yielding the highest R was selected as the “optimal” threshold for distinguishing LB-infected from healthy plant cover in each dataset. Finally, the ratio of LB-infected pixels at the optimal threshold was regressed against plot-level LB severity observations to derive linear regression models for each trial. Although all four datasets were analyzed, only the results at 86 DAP for Trial A and 58 DAP for Trial B are presented in Fig 3C and Fig 3E, respectively, because only a limited number of LB–infected plants were identified during the first assessments in each trial. The same criterion was applied to the second approach as well.

#### Second approach: machine-learning-based regression using multispectral reflectance data

The proposed system was implemented in Python as a modular and customizable ML framework, comprising two interconnected subsystems, as described by [42] (Fig 2B). The first subsystem receives as input the set *u*, which contains L samples *s*_*l*_ ∈ ℝ^5^, for all *l* ∈ {1, 2, … , *L*}, where each *s*_*l*_ represents a five-dimensional multispectral pixel extracted from the polygonal grid of a plot (*u*) in which LB severity observations (*y*) were performed. The input set *u* is then processed through the prediction stage of a K-means clustering algorithm [43], using *K* centroids. Subsequently, a histogram of the input samples *s*_*l*_ is constructed based on the centroids *c*_*k*_ ∈ ℝ^5^, for all *k* ∈ {1, 2, … , *K*}. This procedure yields a histogram vector *x* ∈ ℝ^*K*^, whose elements sum to 1. The K-means model was fitted as described below. Finally, the vector *x* is used as input to the second subsystem, a Gaussian kernel ridge regression model [44,45], which outputs a prediction *y* ∈ ℝ, representing the estimated LB severity of the multispectral five-dimensional pixel set *u*.

#### K-means clustering and histogram-based analysis

To apply the K-means algorithm, this must first be fitted to identify the *K* centroids *c*_*k*_ ∈ ℝ^5^, for all *k* ∈ {1,2, … , *K*}. For this purpose, the M sets *u*_*m*_, for all *m* ∈ {1,2, … , *M*} in the dataset—where M denotes the number of plots—are combined into a single set of elements in ℝ^5^. The K-means algorithm is then applied to this unified set to determine all centroids *c*_*k*_. These centroids are subsequently used in a manner like a bag-of-words system. Each set of pixels corresponding to plot *m* (*u*_*m*_) is encoded using this “alphabet” of *K* centroids, such that each element of *u*_*m*_ is assigned to the nearest centroid to create a frequency histogram. The histogram is normalized so that its elements sum to 1, resulting in a vector *x*_*m*_ ∈ ℝ^*K*^, which represents the coded version of the set *u*_*m*_. This procedure is particularly important because each set *u*_*m*_ may contain a different number of samples (*L*_*m*_ ∈ ℤ); however, the output of this subsystem always returns a fixed-size vector *x*_*m*_of length *K*, independent of the number of pixels in *u*_*m*_. For each dataset (64 and 86 DAP for Trial A, and 38 and 58 DAP for Trial B), an analysis was performed to determine the optimal value of *K* by evaluating values ranging from 32 to 128 in increments of 8. The optimal *K* value was selected as the value that maximizes the R² score during the validation stage of training a Gaussian kernel ridge regression model, as described below.

#### Gaussian kernel ridge regression model

A kernel ridge regression (KRR) model was implemented using a Gaussian kernel (Radial Basis Function) [44,45]. The objective was to find the weight vector (*w*) that minimizes the cost function *L*_*⍺*,*γ*_(*w*), given the hyperparameters *⍺* and *γ*:

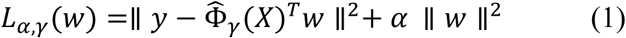

where 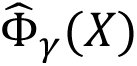 is the vectorial function that generates the kernel matrix 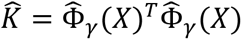, constructed using a radial basis function with parameter *γ*. The vector *y* contains all target values *y*_*m*_, for all *m* ∈ {1,2, … , *M*}, while the matrix *X* includes all input samples 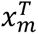, with each sample represented as a row. Each dataset was split into two subsets: 80% for training and validation and 20% for testing. For the training and validation subsets, a 5-fold cross-validation procedure was applied, with four folds used for training and the fifth for validation, yielding five different combinations. After this process, the R^2^ score was calculated for each validation fold, and the mean validation R^2^ was computed for each model. This value was used to optimize α (regularization term), γ (kernel width), and K (K-means) through a grid search that maximized the mean validation R^2^.

### Linear mixed model and rank-based selected coincidence analysis

To perform fair trial-level comparisons among genotypes, a linear mixed model was used to extract the best linear unbiased estimator (BLUE), which represents the adjusted genotype means accounting for trial design and environmental effects. The BLUEs were determined for the raw plot-level AUDPC and LB severity estimated using normalized difference vegetation index-based linear regression (LB_LR_) and kernel ridge regression-based machine-learning models (LB_KRR_). Thus, separate linear fixed models were fitted for Trials A and B, using the “statgenSTA” R package [46].

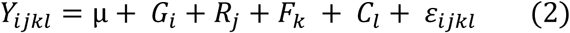

where *Y*_*ijkl*_ is the performance of the i^th^ genotype in the j^th^ replication located in the k^th^ row and l^th^ column. µ is the overall mean, *G*_*i*_ is the fixed effect of the i^th^ genotype, while *R*_*j*_ (j^th^ replication effect), *F*_*k*_ (k^th^ row effect), and *C*_*l*_ (l^th^ column effect) were treated as random effects to account for spatial variability. *ɛ*_*ijkl*_ is the error term associated with each *Y*_*ijkl*_.

Later, Spearman’s rank correlation was used to assess the relationship between BLUEs of AUDPC and BLUEs of estimated LB severity (LB_LR_ and LB_KRR_). Kernel density distributions of standardized BLUE values (z-scores) were computed using the “stats” package in R to visualize and compare distribution patterns between AUDPC and model-estimated LB severity. Furthermore, the percentage of selected coincidence (SC) between AUDPC- and LB-based rankings within the top 10%, 20%, and 30% of clones ranked by AUDPC was calculated comparing the selections from both rankings. This analysis was conducted for both UAV assessment dates to determine whether a single time-point estimate of LB severity could provide comparable genotype rankings to those obtained with the integrative AUDPC indicator for early breeding selection.

### Model performance evaluation

Model performance was evaluated by comparing observed and estimated LB severity values using R^2^ and the Root Mean Square Error (RMSE). All statistical analyses were performed using the R programming language [47].

## Results

### Estimation of late blight severity using vegetation indices

NDVI was the most representative vegetation index for assessing plot-level LB severity, exhibiting the strongest correlation with LB severity across almost all datasets (−0.55 to −0.86, p < 0.001), except at 38 DAP in Trial B (Table 1). GNDVI and RE ranked second in Trials A and B, respectively, across both assessment dates (Table 1). Overall, plot-level NDVI showed stronger correlations with LB severity in the second assessment (−0.41 to −0.86) than in the first assessment (−0.18 to −0.55) (Table 1). Sweeping NDVI thresholds produced bell-shaped response curves relating NDVI threshold values to the correlations between threshold-based LB severity estimates and observed ones, with correlation values ranging from 0.21 to 0.76 in Trial A and from 0.35 to 0.90 in Trial B (Fig 3A). The strongest correlations were found at NDVI thresholds of 0.60 for Trial A and 0.55 for Trial B, both during the second assessment (Fig 3A). Linear regression between genotype-level observed LB severity and severity estimated using optimal NDVI thresholds showed contrasting performances across trials and assessment dates (Fig 3C, E). In Trial A, model performance was weak at 64 DAP (R^2^ = 0.19, p < 0.001; RMSE = 15.5%) but improved at 86 DAP (R^2^ = 0.65, p < 0.001; RMSE = 39.2%). In Trial B, the model performance showed a similar pattern, being weak at 38 DAP (R^2^ = 0.23, p < 0.001; RMSE = 47.8) and substantially stronger at 58 DAP (R^2^ = 0.89, p < 0.001; RMSE = 11.9%).

**Table 1.** Pearson correlation (R) values between mean vegetation indexes—Normalized Difference Vegetation Index (NDVI), Green Normalized Difference Vegetation Index (GNDVI), Red Edge reflected (RE), and Normalized Difference Red Edge (NDRE)—and Late Blight severity observations from Trials A (1,476 plots) and B (5,490 plots). Asterisks (**\***) indicate statistically significant correlations at p < 0.001. DAP = Days after planting.

| Vegetation Index | Trial A |  | Trial B |  |
| --- | --- | --- | --- | --- |
|  | 64 DAP | 86 DAP | 38 DAP | 58 DAP |
| NDVI | **-0.55 \*** | **-0.86 \*** | -0.33 * | **-0.60 \*** |
| GNDVI | -0.51 * | -0.64 * | -0.22 * | -0.41 * |
| RE | -0.40 * | -0.57 * | **-0.37 \*** | -0.46 * |
| NDRE | -0.50 * | -0.54 * | -0.18 * | -0.45 * |

### Estimation of LB severity from multispectral reflectance data using clustering and KRR

The optimal number of clusters (*K*) varied across trials and assessment dates (Table 2, Fig 3B). In Trial A, the highest performance was achieved with *K* = 96 at 64 DAP and *K* = 88 at 86 DAP, whereas in Trial B, the highest performance was obtained with *K* = 104 at 38 DAP and *K* = 96 at 58 DAP (Table 2, Fig 3B). Overall, KRR models showed stronger predictive performance at the second assessment date, characterized by smaller values of the hyperparameters α and without pattern in the hyperparameter γ (Table 2). In Trial A, the model performed better at 86 DAP (R² = 0.80, p < 0.001; RMSE = 6.6%) than at 64 DAP (R² = 0.46, p < 0.001; RMSE = 4.8%). In Trial B, the model performance was lower at 38 DAP (R² = 0.46, p < 0.001; RMSE = 5.6%) and higher at 58 DAP (R² = 0.88, p < 0.001; RMSE = 7.9%) (Table 2, Fig 3D, F).

**Table 2.**
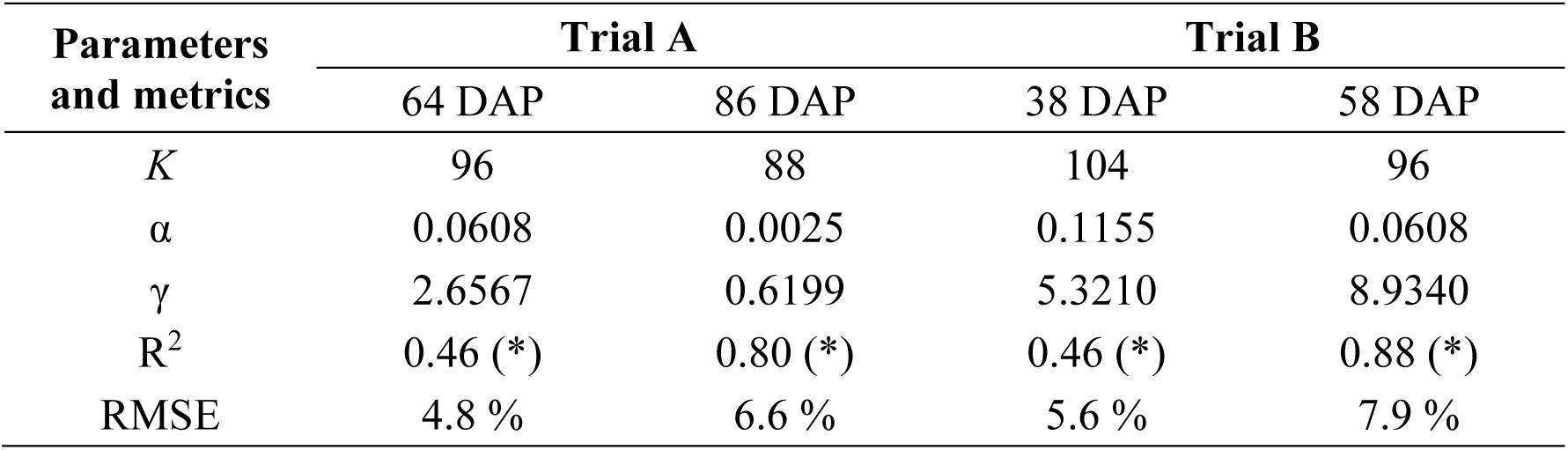
Tuned hyperparameters—number of centroids (*K*), regularization term (α), and kernel width (γ)—of the combined K-means clustering and Kernel Ridge Regression model, and regression metrics—coefficient of determination (R^2^) and root mean square error (RMSE)—comparing model-estimated and observed late blight severity in two large potato breeding trials. Asterisks (*) indicate statistically significant correlations at p < 0.001. DAP = Days after planting.

### Relationship between AUDPC and estimated LB severity

A wide range of disease susceptibility levels was observed across genotypes in both trials, with BLUEs of AUDPC ranging from 24.1 to 2007.0%-days in Trial A and from 32.2 to 2322.8%-days in Trial B (Fig 4). The kernel density distributions of BLUEs of LB_LR_ and LB_KRR_ generally followed the AUDPC distribution across trials, although model-based estimates showed narrower spreads and slight shifts relative to AUDPC, particularly in the second assessment of Trial B (Fig S3). During the first UAV assessment, BLUEs of model-based LB severity estimates were relatively low for most genotypes, particularly in Trial A, resulting in a weaker association with AUDPC (Spearman’s coefficient ranging from 0.311 to 0.707) (Fig S2). In the second UAV assessment, BLUEs of LB severity showed a more homogeneous distribution across low, intermediate, and high values, resulting in stronger correlations with AUDPC (Spearman’s coefficient ranging from 0.774 to 0.895) (Fig 4). The percentage of SC between AUDPC- and LB-based rankings was lower during the first UAV assessment, ranging from 17.1% to 48.1% in Trial A and from 46.8% to 70.5% in Trial B, depending on the selection intensity and LB estimation method (Fig S2). In contrast, substantially higher SC values were observed in the second UAV assessment, ranging from 47.6% to 77.4% in Trial A and from 80.9% to 91.5% in Trial B (Fig 4). Overall, LB estimates obtained with KRR showed better performance in selecting top-ranked genotypes across assessment dates and selection intensities (Figs S2 and 4).

**Fig 4.**
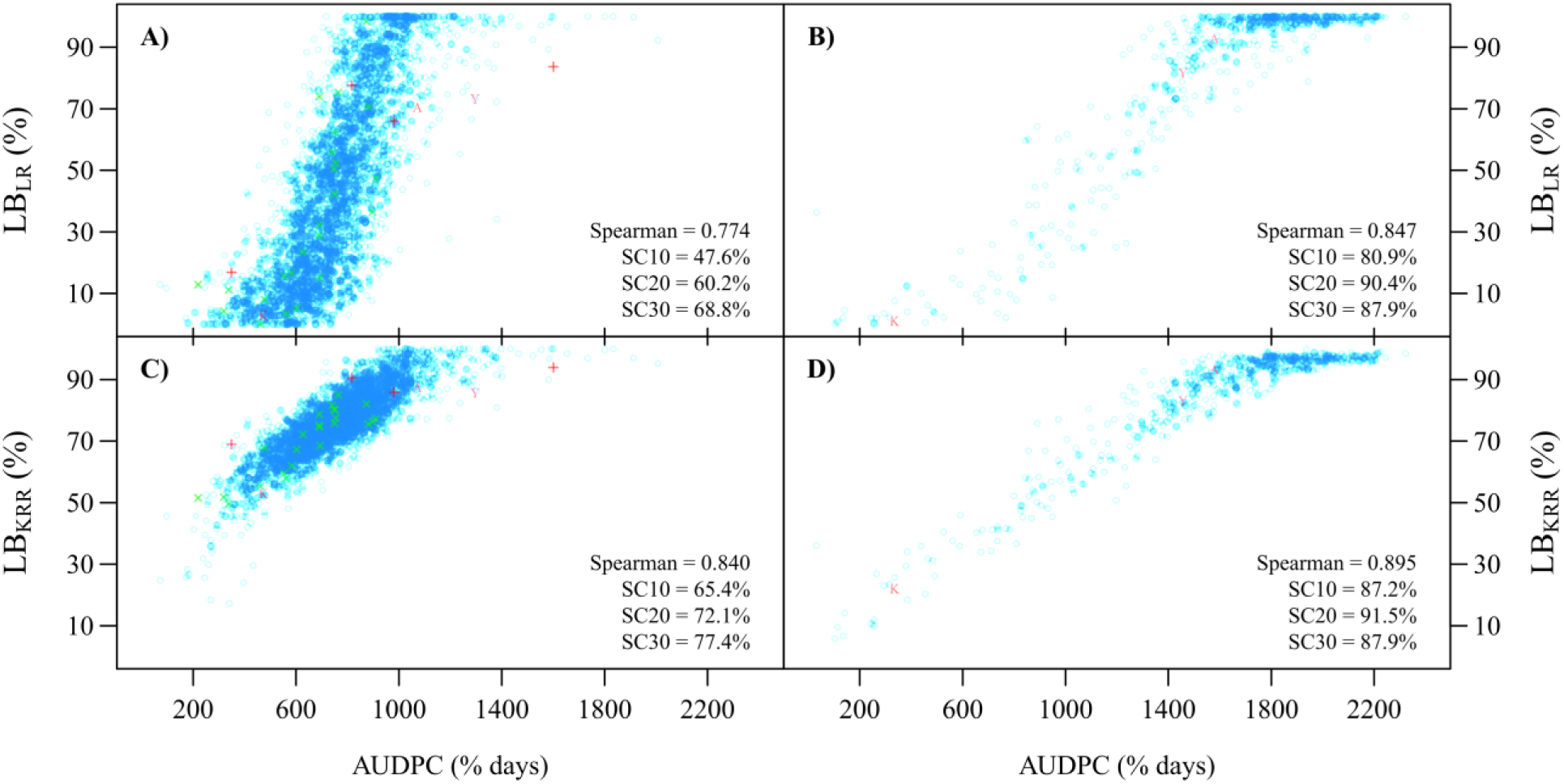
BLUEs of AUDPC vs. UAV-based LB severity estimates. Best linear unbiased estimators (BLUEs) of the area under the disease progress curve (AUDPC) versus BLUEs of estimated late blight (LB) severity in large potato breeding trials. LB severity was estimated by two approaches: normalized difference vegetation index-based linear regression (LB_LR_; panels (A) and (B)) and kernel ridge regression-based machine-learning models (LB_KRR_; panels (C) and (D)). Panels (A) and (C) show results for 2,745 clones (Trial A) evaluated at 86 days after planting (DAP), whereas panels (B) and (D) show 492 accessions (Trial B) evaluated at 58 DAP. Red symbols (with A=Amarilis, K=Kori, and Y=Yungay common to both trials) and green symbols represent check and parent varieties, respectively. SC10, SC20, and SC30 denote the percentage of selected coincidence between AUDPC- and LB-based rankings within the top 10%, 20%, and 30% of clones ranked by AUDPC.

Among the checks cultivars, Amarilis and Yungay consistently exhibited high BLUEs of AUDPC across both trials, with values ranging from 1,071.7 to 1,296.1 % -days in Trial A and from 1,456.1 to 1,578.2 % -days in Trial B. Consistent with these high disease progress values, both cultivars also showed elevated BLUEs of LB severity, as estimated from NDVI (70.5 to 73.2 % in Trial A and 81.7 to 92.8 % in Trial B) and KRR-based models (85.7-87.4 % in Trial A and 83.1-90.3 % in Trial B), in agreement with their known susceptibility profiles. In contrast, the resistant cultivar Kory consistently displayed low BLUEs of AUDPC in both trials, with values of 473.7 %-days in Trial A and 336.0 %-days in Trial B. Correspondingly, BLUEs of LB severity were markedly lower, ranging from 2.8% (NDVI-based) and 53.0% (KRR-based) in Trial A to 1.0% (NDVI-based) and 22.0% (KRR-based) in Trial B (Fig 4), reinforcing its comparatively resistant response.

## Discussion

### UAV-based remote sensing combined with K-means-KRR improves late blight severity estimation over NDVI-based models

Late blight progresses through two distinct phases: an initial biotrophic phase involving physiological changes, mainly chlorophyll reduction [48–50], without visible symptoms [51], and a subsequent necrotrophic phase characterized by structural tissue damage, including cell wall degradation [52,53] and membrane disruption [54–57]. These biochemical and structural modifications directly alter the canopy’s spectral reflectance properties, making remote sensing a viable approach for detecting and quantifying disease severity. Spectral reflectance in the near-infrared (NIR) region is primarily governed by the internal leaf structure and dry matter content, including proteins, lignin, and cellulose [58,59], whereas chlorophyll degradation predominantly affects reflectance in the blue and red absorption bands [38,60]. Consequently, vegetation indices that combine NIR with blue or red spectral bands, such as NDVI, enhance the remote detection of LB symptoms across both biotrophic (early) and necrotrophic (advanced) stages. Consistent with this principle, NDVI showed strong associations with LB severity in this study. The identification of threshold values (0.6 and 0.55) enabled objective classification of healthy and diseased canopy tissues across assessment periods. Similar threshold-based approaches have proven effective for disease severity detection in several crops [61–64], supporting the robustness of vegetation index-derived metrics for plant disease monitoring.

Beyond vegetation indices, integrating K-means clustering with Kernel Ridge Regression provided a scalable framework for estimating LB severity across large breeding trials. Clustering allowed heterogeneous pixel distributions to be summarized into fixed-length representations, facilitating high-throughput phenotyping across variable canopy structures [65]. The KRR-based model for estimating LB severity (LB-KRR) further captured nonlinear relationships within multispectral reflectance data, leading to consistently improved predictive performance relative to NDVI-based linear models [66,67]. Fine-tuning key hyperparameters enhanced the model’s sensitivity to both early physiological changes and severe late-stage damage. As a result, the LB-KRR approach achieved superior predictive performance during early assessments in both trials, whereas at later stages, improvements were primarily reflected in reduced RMSE, indicating enhanced plot-level prediction precision.

### Robust ML performance under breeding-scale conditions

A key contribution of this study lies in the scale at which ML–based LB severity estimation was evaluated. While previous studies using UAV imagery and advanced ML or deep learning models have reported high predictive performance, they have generally been limited to small experimental populations (typically fewer than 15 genotypes) [17,25,68,69] constraining their transferability to breeding-scale applications. In contrast, the present study evaluated ML–based disease severity estimation across large, genetically diverse populations comprising 2,745 clones in Trial A and 492 accessions in Trial B. This scale introduces substantial variability—such as heterogeneous canopy architectures, differential disease progression, and variable spectral responses across genotypes—that is rarely addressed in literature. Despite these challenges, the K-means–KRR framework maintained robust predictive performance, highlighting its scalability and suitability for early-generation breeding pipelines characterized by large populations and high phenotypic diversity.

### UAV-based methods can complement and, in specific contexts, partially replace repeated visual assessments

Conducting UAV-based phenotyping across multi-environment breeding trials introduces logistical constraints that can affect both data availability and model performance. Coordinating flights across locations requires suitable weather conditions, trained personnel, and available equipment, which often limit the number and timing of image acquisitions during the growing season. Similar challenges have been reported in large-scale UAV phenotyping studies [70–72]. Nevertheless, in breeding programs screening hundreds or thousands of genotypes, the primary objective is often to achieve high selection accuracy through reliable ranking of genotypes to prioritize promising clones for subsequent evaluation stages [73], rather than to obtain a precise quantification of LB severity. In this context, defining the optimal number and timing of UAV acquisitions becomes critical. Disease progression, canopy development, and environmental conditions strongly influence the spectral expression of plant stress and, in turn, the detectability of disease using optical sensors [62,74]. Evaluating the contribution of individual UAV acquisitions across environments and disease stages is therefore essential to identify the most informative phenotyping windows. In this study, UAV flights conducted during intermediate to advanced stages of LB disease progression (e.g., the second flight) provided the most relevant information for genotype ranking and selection decisions. The high coincidence observed for the top 10% of genotypes (SC10 = 65.4% in Trial A and 87.2% in Trial B with KRR) indicates that a limited number of acquisitions—and potentially even a single well-timed flight—can approximate AUDPC-based selections while reducing the need for repeated field assessments.

Early-generation trials typically involve large populations, limited replication, and high phenotypic variability, making precise manual assessments challenging [73]. Given these conditions, the predictive performance observed in this study—although variable across environments and assessment dates—appears sufficient to support early-stage selection by enabling the identification of highly susceptible genotypes and prioritizing promising clones for further evaluation. Consistent with previous high-throughput phenotyping studies, even moderate prediction accuracies can contribute to genetic gain by enabling the screening of much larger breeding populations [75]. Furthermore, because visual assessments are often subject to observer bias and inconsistency across environments, UAV-derived estimates—even when moderately correlated—can improve the objectivity and efficiency of selection. When integrated with efficient data-processing pipelines, these UAV-derived phenotypes can provide timely and objective information that complements genomic-assisted breeding strategies and enhances selection efficiency in large-scale breeding programs [76].

## Conclusion

UAV-based multispectral imaging provides a viable approach for estimating late blight severity under breeding field conditions, capturing biologically meaningful variation across large and genetically diverse populations, including 2,745 clones and 492 accessions. While NDVI offers a simple and effective baseline for disease detection, the integration of K-means clustering and kernel-based regression improves the representation of spectral–disease relationships by capturing nonlinear patterns, resulting in more accurate severity estimation and genotype ranking. When expressed as BLUEs, UAV-derived estimates showed strong and biologically consistent relationships with AUDPC and achieved higher selection coincidence, particularly at intermediate to advanced stages of disease development. These findings indicate that image-based phenotyping can reliably support genotype ranking for selection, especially when acquisitions are conducted at appropriate disease stages. Overall, UAV-based phenotyping represents a scalable and practical complement to conventional late blight assessments in early-generation breeding trials, where large population sizes limit precise manual evaluation. Future work should focus on optimizing acquisition strategies and evaluating their impact on selection decisions across diverse environments.

## Supporting information

**S1 Fig. Weather conditions during potato trials and UAV acquisitions**. Weather conditions in the study area of Oxapampa, Peru (September 2021–February 2022). Minimum (blue lines) and maximum (red lines) temperatures are on the left y-axis, while precipitation (PP, cyan bars) and relative humidity (RH, black dotted lines) are on the right. Vertical green and orange dashed lines mark planting and harvest for the first (A) and second (B) trials. Red arrows indicate the two unmanned aerial vehicle flight dates for multispectral imagery acquisition.

**S2 Fig. AUDPC vs. NDVI- and KRR-based LB severity: implications for selection**. Best linear unbiased estimators (BLUEs) of the area under the disease progress curve (AUDPC) versus BLUEs of estimated late blight (LB) severity in large potato breeding trials. LB severity was estimated by two approaches: normalized difference vegetation index-based linear regression (LB_LR_; panels (A) and (B)) and kernel ridge regression-based machine-learning models (LB_KRR_; panels (C) and (D)). Panels (A) and (C) show results for 2,745 clones (Trial A) evaluated at 64 days after planting (DAP), whereas panels (B) and (D) show 492 accessions (Trial B) evaluated at 38 DAP. Red symbols (with A=Amarilis, K=Kori, and Y=Yungay common to both trials) and green symbols represent check and parent varieties, respectively. SC10, SC20, and SC30 denote the percentage of selected coincidence between AUDPC- and LB-based rankings within the top 10%, 20%, and 30% of clones ranked by AUDPC.

**S3 Fig. Density distributions of AUDPC and LB severity estimates.** Kernel density distributions of standardized values (z-scores) of best linear unbiased estimators (BLUEs) of the area under the disease progress curve of late blight (AUDPC; light blue) and the BLUEs of late blight severity estimated using normalized difference vegetation index-based linear regression (LB_LR_; green) and kernel ridge regression-based machine-learning models (LB_KRR_; pink). Panels (A) and (C) show results for 2,745 clones (Trial A) evaluated at 64 and 86 Days After Planting (DAP), and panels (B) and (D) for 492 accessions (Trial B) evaluated at 38 and 58 DAP.

## Notes

https://data.cipotato.org/dataset.xhtml?persistentId=doi:10.21223/YT4UZU

https://data.cipotato.org/dataset.xhtml?persistentId=doi:10.21223/GCJRH6

### Competing Interest Statement

The authors have declared no competing interest.

### Funding

- This research was supported, in whole or in part, by the Bill & Melinda Gates Foundation (grant number OPP1070785); the CGIAR Research Program on Roots, Tubers, and Bananas (RTB); the One CGIAR Initiative on Plant Health; and the CGIAR Sustainable Farming Science Program and the Breeding for Tomorrow Science Program. CGIAR is a global research partnership dedicated to transforming food, land, and water systems in the face of the climate crisis and advancing a food-secure future.: OPP1070785

