## Supplementary figures and images for "Prediction of late blight severity in a large panel of potato genotypes using low-altitude aerial images and machine learning methods"

### Supplementary Figure 1 (S1 Fig)

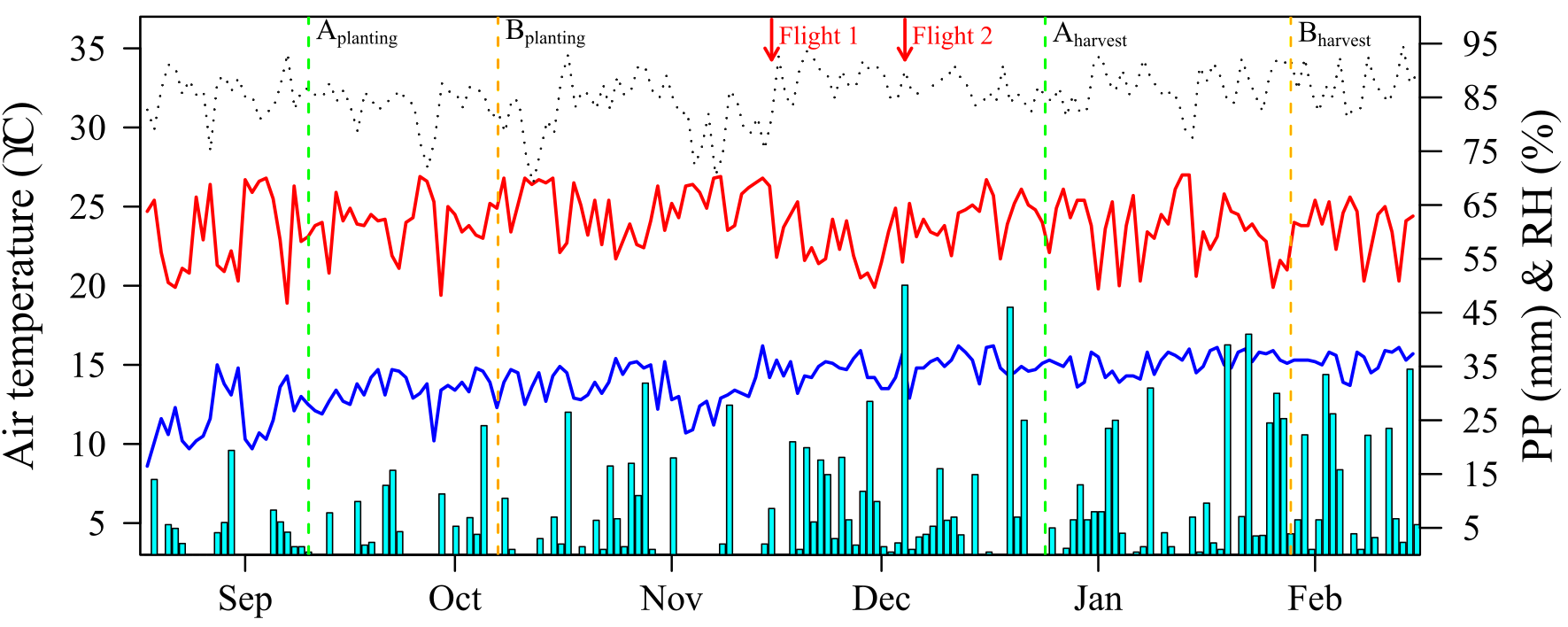

### Supplementary Figure 2 (S2 Fig)

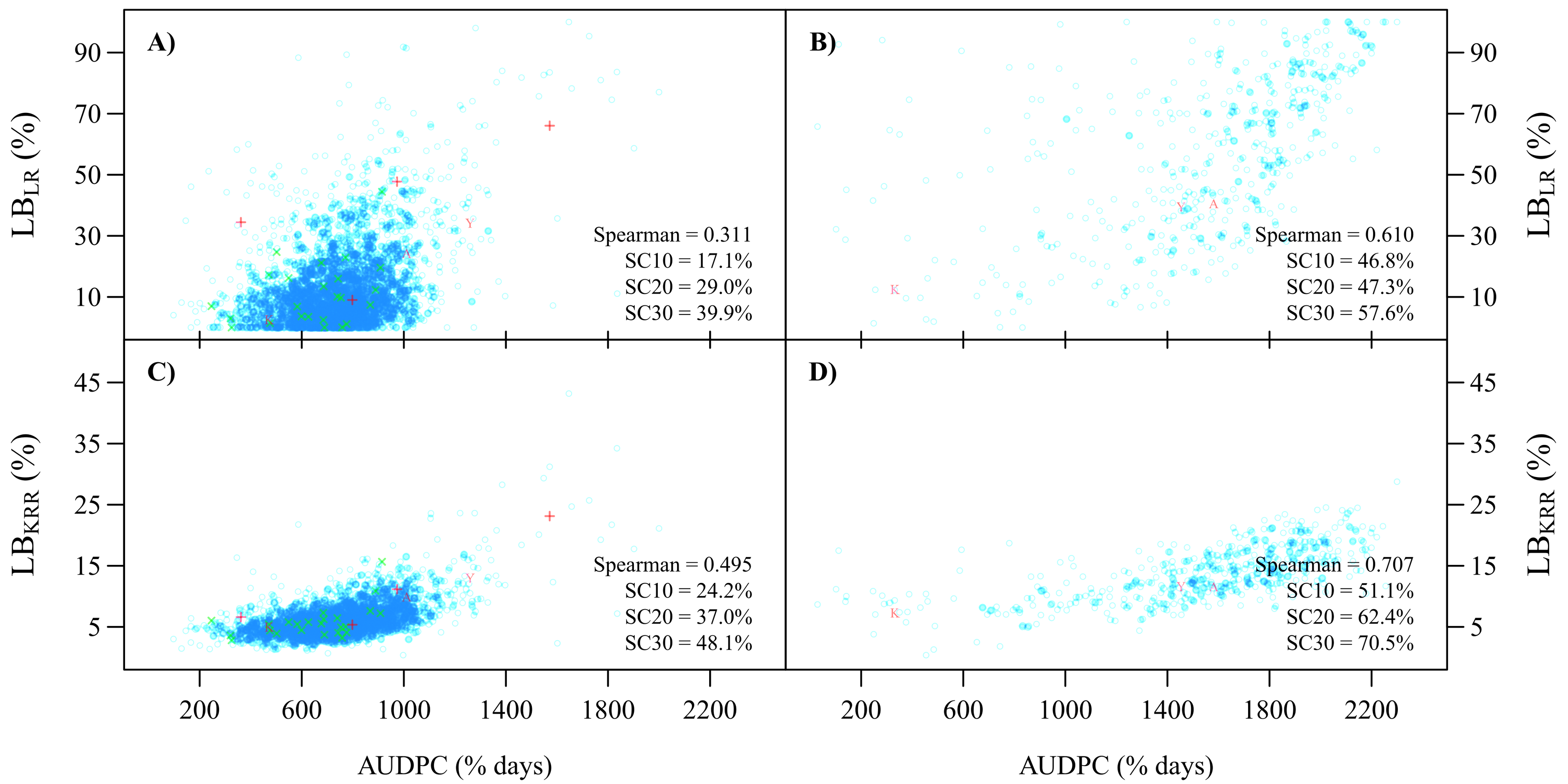

### Supplementary Figure 3 (S3 Fig)

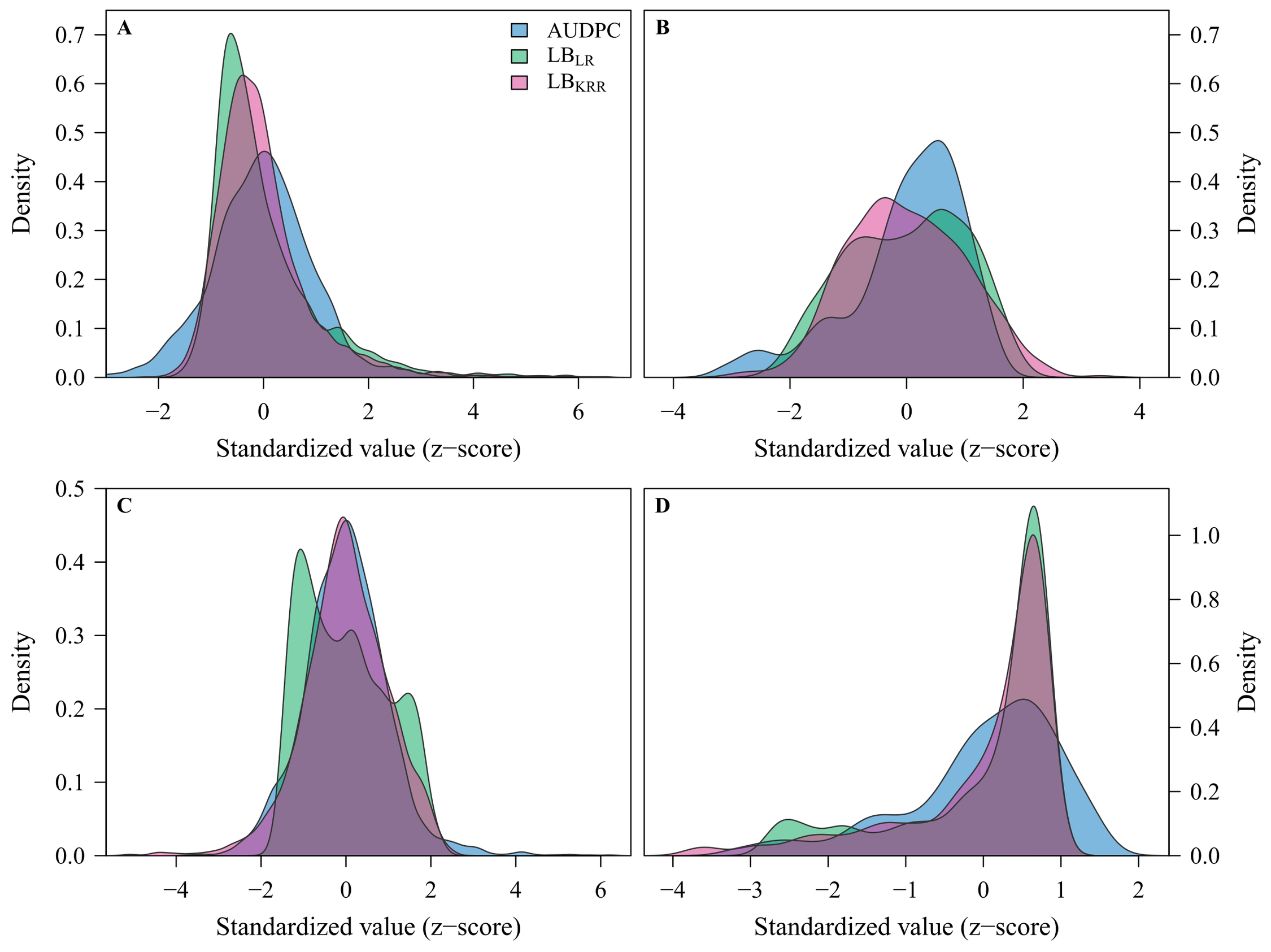
